# Assessment of pupillometry across different commercial systems of laying hens to validate its potential as an objective indicator of welfare

**DOI:** 10.1101/2025.06.06.658220

**Authors:** Elyse Mosco, David Kilroy, Arun HS Kumar

## Abstract

**Background:** Reliable and non-invasive methods for assessing welfare in poultry are essential for improving evidence-based welfare monitoring and advancing management practices in commercial production systems. The iris-to-pupil (IP) ratio, previously validated by our group in primates and cattle, reflects autonomic nervous system balance and may serve as a physiological indicator of stress in laying hens. This study evaluated the utility of the IP ratio under field conditions across diverse commercial layer housing systems.

**Materials and Methods:** 296 laying hens (Lohmann Brown, n = 269; White Leghorn, n = 27) were studied across four locations in Canada housed under different systems: Guelph (indoor; pen), Spring Island (outdoor and scratch; organic), Ottawa (outdoor, indoor and scratch; free-range), and Toronto (outdoor and hobby; free-range). High-resolution photographs of the eye were taken under ambient lighting. Light intensity was measured using the light meter app. The IP ratio was calculated using NIH ImageJ software. Statistical analysis included one-way ANOVA and linear regression using GraphPad Prism.

**Results:** Birds housed outdoors had the highest IP ratios, followed by those in scratch systems, while indoor and pen-housed birds had the lowest IP ratios (p < 0.001). Subgroup analyses of birds in Ottawa and Spring Island farms confirmed significantly higher IP ratios in outdoor environments compared to indoor and scratch systems (p < 0.001). IP ratio correlated weakly with ambient light intensity (r = 0.25) and age (r = 0.05), indicating minimal influence of these variables. Although White Leghorn hens showed lower IP ratios than Lohmann Browns, this difference was confounded by housing type; all White Leghorns were housed in pens. Thus, housing system but not breed was the primary driver of IP variation.

**Conclusion:** IP ratio is a robust, non-invasive physiological marker of welfare assessment in laying hens, sensitive to housing environment but minimally influenced by light or age. Its potential for integration with digital imaging technologies supports its use in scalable welfare assessment protocols.

## Background

Welfare assessment in animals has increasingly emphasized the need for objective, non-invasive measures that reliably reflect physiological and psychological states [1-4]. In this context, pupillometry (the measurement of pupil dynamics) has gained traction as a valuable tool, particularly in human psychological and neurological research [5,6]. The pupil’s diameter is known to change in response to a variety of internal and external stimuli, including emotional arousal, cognitive effort, and stress [7,8]. These changes are governed by the autonomic nervous system: the radial dilator muscles of the iris are under sympathetic control, while the circular sphincter muscles are regulated by the parasympathetic system [7,8]. Under ambient light conditions, fluctuations in pupil size are predominantly driven by shifts in this autonomic balance, making pupillary dynamics a promising biomarker of stress, welfare and arousal [9,10]

Current approaches to measuring welfare in poultry encompass a combination of behavioral, physiological, and environmental assessments. Traditionally, welfare evaluation relies heavily on behavioral indicators such as feather pecking, aggression, and the expression of natural behaviors like dust bathing, perching, and foraging [11-13]. These behaviors provide valuable insight but can be subjective and influenced by observer bias. Physiological markers, including corticosterone levels in blood or feathers, heterophil-to-lymphocyte ratios, and stress hormone assays, offer more objective data but are often invasive, requiring handling or sampling that can itself induce stress. Environmental assessments, such as litter quality, space allowance, and air quality, are also used to predict welfare conditions indirectly. Additionally, scoring systems and welfare audits such as the Welfare Quality® assessment protocol combine these parameters into standardized frameworks [11-15]. However, these methods can be time-consuming, labor-intensive, and inconsistent across different production systems. There is a growing need for more efficient, objective, and non-invasive tools that can reliably assess welfare on-farm without disrupting normal animal behavior [11,15]. Emerging technologies, including automated behavior tracking, thermal imaging, and physiological metrics like pupillometry, are now being explored to complement or potentially replace traditional welfare assessment techniques, offering a promising avenue for real-time, technology-driven monitoring of welfare in poultry husbandry.

In birds, including the domestic hen (*Gallus gallus domesticus*), the pupil has been shown to respond sensitively to meaningful stimuli [16,17]. Previous studies have demonstrated that hens exhibit pupil dilation in response to aversive or arousing stimuli, suggesting that pupil measurements may serve as a quantitative indicator of welfare-relevant experiences [17-20]. Despite these findings, pupillometry remains underexplored in avian species as a standardized welfare assessment tool. The current study builds on our previous research in primates and cattle [7,8], where we reported the iris-to-pupil (IP) ratio as an objective, non-invasive indicator of stress and welfare status. The IP ratio reflects the proportional relationship between the iris and the pupil, offering a dimensionless metric that is less sensitive to absolute size and more reflective of autonomic nervous system activity. This study aimed to evaluate the relevance and reliability of the IP ratio in laying hens across various commercial rearing systems, including organic, free-range, and loose-caged environments. A specific focus of this investigation was to determine whether pupil parameters, particularly the IP ratio, are affected by ambient light intensity and whether they vary in association with stress-inducing or stress-relieving aspects of the housing environment. Given that light is a key environmental variable influencing pupil size, controlling or accounting for its effects is critical for the interpretation of pupillometric data in field conditions [21-23]. Thus, light intensity was recorded in all settings to assess its potential impact on IP ratio measurements. We hypothesized that pupil parameters, specifically the IP ratio, will differ significantly between husbandry systems due to varying degrees of environmental enrichment and associated welfare status. By integrating pupillometry into field-based welfare assessments, this study aimed to advance our understanding of how housing systems affect the physiological well-being of laying hens and to explore the potential of IP ratio as a scalable, digital imaging–based welfare metric in poultry husbandry and perhaps other avian species.

## Materials and Methods

This study was designed to assess variations in pupillometry in laying hens housed in different commercial systems, i.e., Hobby free range, free-range, organic (EU-compliant), and loose caged. Laying hens from these systems were photographed and analyzed to assess variation in pupil size, specifically the iris-to-pupil (IP) ratio, which we have previously identified as an objective measure of welfare in primates and cattle [7,8]. Building on this prior work, the current study aimed to evaluate the relevance and applicability of the IP ratio as a non-invasive welfare indicator in commercial laying hens. As this study employed a fully non-invasive approach, involving only the photographic capture of eye images without any handling, restraint, or physical interaction with the birds, it was deemed to pose minimal risk to animal welfare. Consequently, the study was exempted from full institutional animal ethics review in accordance with the guidelines governing observational research involving animals. Nonetheless, all procedures were conducted in alignment with established ethical standards to ensure the comfort and well-being of the hens throughout the data collection process following informed consent from the facility owners.

Data were collected from four locations in Canada and a total of 296 laying hens were included in this study, representing a range of commercial production systems and environmental conditions. The housing environments included outdoor, indoor, scratch areas and pens across different system types, i.e., hobby free-range (6 birds), free-range (169 birds), organic EU-compliant (94 birds), and loose caged (indoor pen-based housing) (27 birds). The organic system was represented by Olive Farm on Spring Island, British Columbia, where hens were kept outdoors with access to a scratch area. The free-range system was studied at two sites: Slevin Farm in Toronto, Ontario, a small-scale hobby farm with outdoor access and Kelly Farm in Ottawa, Ontario, which included both indoor and outdoor access along with a scratch area. The loose caged system was represented by a controlled floor pen facility at a commercial farm in Guelph, Ontario, which exclusively housed White Leghorn hens. Across these sites, the housing environments included outdoor access, indoor housing, scratch areas (cage-free enriched), and enclosed pens.

A subgroup analysis was conducted using data from birds at Kelly and Olive farms to assess differences in the IP ratio across three housing sub-environments, i.e., outdoor, indoor, and scratch areas. At Kelly Farm, which followed a free-range system, birds were photographed in three distinct environments: outdoor (n = 60), indoor (n = 38), and scratch area (n = 71). At Olive Farm, managed under an organic EU-compliant system, birds were similarly photographed in outdoor (n = 42) and scratch (n = 52) settings. This subgrouping allowed for a targeted comparison of IP ratios among birds exposed to different levels of environmental enrichment and access within the same farm, helping to control for variability related to location and management practices. By comparing these subgroups, the study aimed to determine whether specific aspects of the housing environment such as outdoor access versus enriched indoor spaces had a measurable impact on physiological stress, as reflected by changes in the IP ratio.

Two commercial breeds of laying hens were included in the study: Lohmann Brown (n = 269) and White Leghorn (n = 27). Hens were sampled randomly within each system, ensuring adequate representation of each breed where available. Photographs of the hens’ eyes were taken using a high-resolution digital camera in the birds’ natural housing environments to minimize handling-induced stress that could alter pupil size. Each photograph was taken under existing ambient lighting conditions, and light intensity (in lux) was measured using a light meter application on an iPhone. Measurements were taken by holding the device with the forward-facing camera oriented parallel to the floor and directed toward the primary light source, positioned at the level of the birds’ eyes. This method allowed for consistent and practical estimation of ambient light intensity across different sites and housing systems.

Digital image analysis was conducted using NIH ImageJ software. The iris and pupil area were measured in pixels from each photograph (Figure 1A), and the iris-to-pupil area ratio (IPR) was calculated for normalization. This ratio served as the primary metric for comparing pupil size across housing systems, as it controls for differences in eye size and camera distance. One observer, blinded to housing conditions, performed the image measurements to reduce bias.

**Figure 1.**
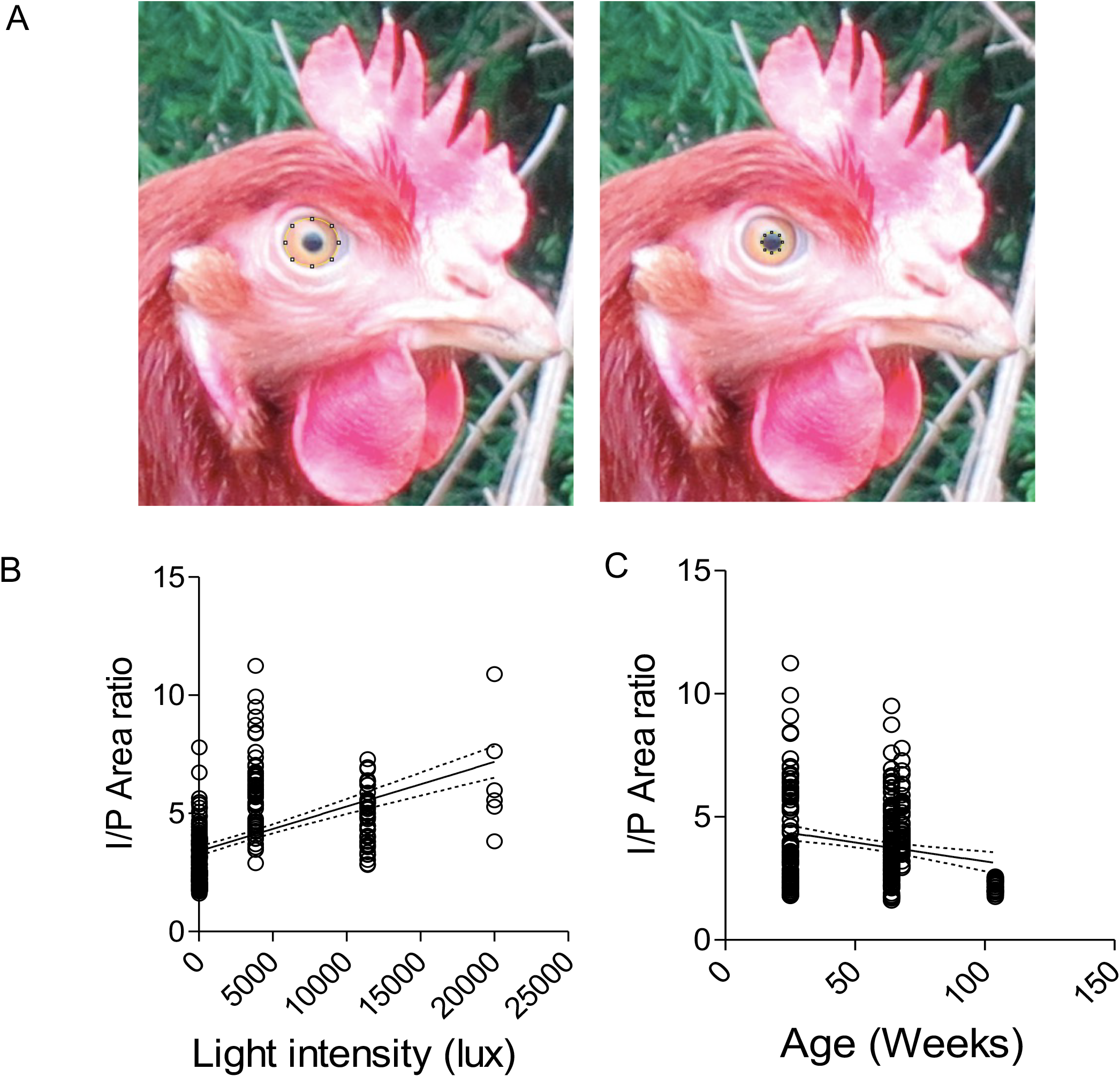
A: Representative image of Layer eye to show the method used to measure iris (left) and pupil (right) area using Image-J software. B: Linear regression analysis between Iris-Pupil (I/P) area ratio (1.6 to 11.2) and light intensity (21 to 20000 Lux). C: Linear regression analysis between Iris-Pupil (I/P) area ratio (1.6 to 11.2) and age (22 to 104 Weeks). Regression line along with it 95% confidence interval is shown (n = 296).

Statistical analyses were performed using GraphPad Prism, version 5. One-way analysis of variance (ANOVA, followed post hoc Bonferroni’s Multiple Comparison Test) or Student t-test was used to determine significant differences in IP ratios among different groups. In addition, linear regression analysis was performed to evaluate the relationship between ambient light intensity/age and pupil diameter. Statistical significance was set at p < 0.05, and all results were reported as mean ± standard deviation of all data collected.

## Results

We compared the relationship between light intensity or age of birds and IP ratio among our entire cohorts of layers. The correlation coefficient between light intensity and age with IP ratio was 0.25 and 0.05 respectively, suggesting a weak correlation (Figure 1B). This very low correlation coefficient, especially with a light intensity rage of 21 to 20000 lux, suggests minimal influence of ambient light variations on pupillary response in birds (Figure 1B). The correlation between age of birds ranging from 25 to 104 weeks and IP ratio although weaker showed a regression line with negative slope (Figure 1C). These data validate the reliability of IP ratio measurement as an objective measure of autonomic balance (sympathetic and parasympathetic system) under field settings where variations in ambient light conditions are normally observed.

Having established the minimal influence of ambient light variations on IP ratio, we next assessed the influence of different layer husbandry systems on IP ratio. The average of IP ratio of all birds (n=296) in the study cohorts was 3.93±1.76, while the average of IP ratio of birds in the hobby FR, free range, organic and loose cage system was 6.52±2.47, 3.88±1.91, 4.37±1.16 and 2.9±0.24 respectively. The IP ratio of birds under hobby free range system was 68.23 %, 49.22% and 198.16% greater than the IP ratio of birds in free range, organic and loose cage systems respectively. The average of IP ratio of all birds in the study cohorts was significantly (p < 0.01) lower than that of birds from hobby FR system but was significantly (p < 0.001) greater than that of birds from loose cage system (Figure 2). The IP ratio of birds from hobby FR system was significantly greater than the IP ratio of birds from free range (p > 0.01), organic (p > 0.05) and loose cage systems (p < 0.001) (Figure 2). The IP ratio of birds from loose cage system was significantly (p < 0.001) lower than the IP ratio of birds from free range and organic systems (Figure 2). The IP ratio of birds in hobby FR system was greater than the population average, whereas the IP ratio of birds in free range and organic systems was similar to the population average while the IP ratio of birds in loose cage system was smaller than the population average.

**Figure 2.**
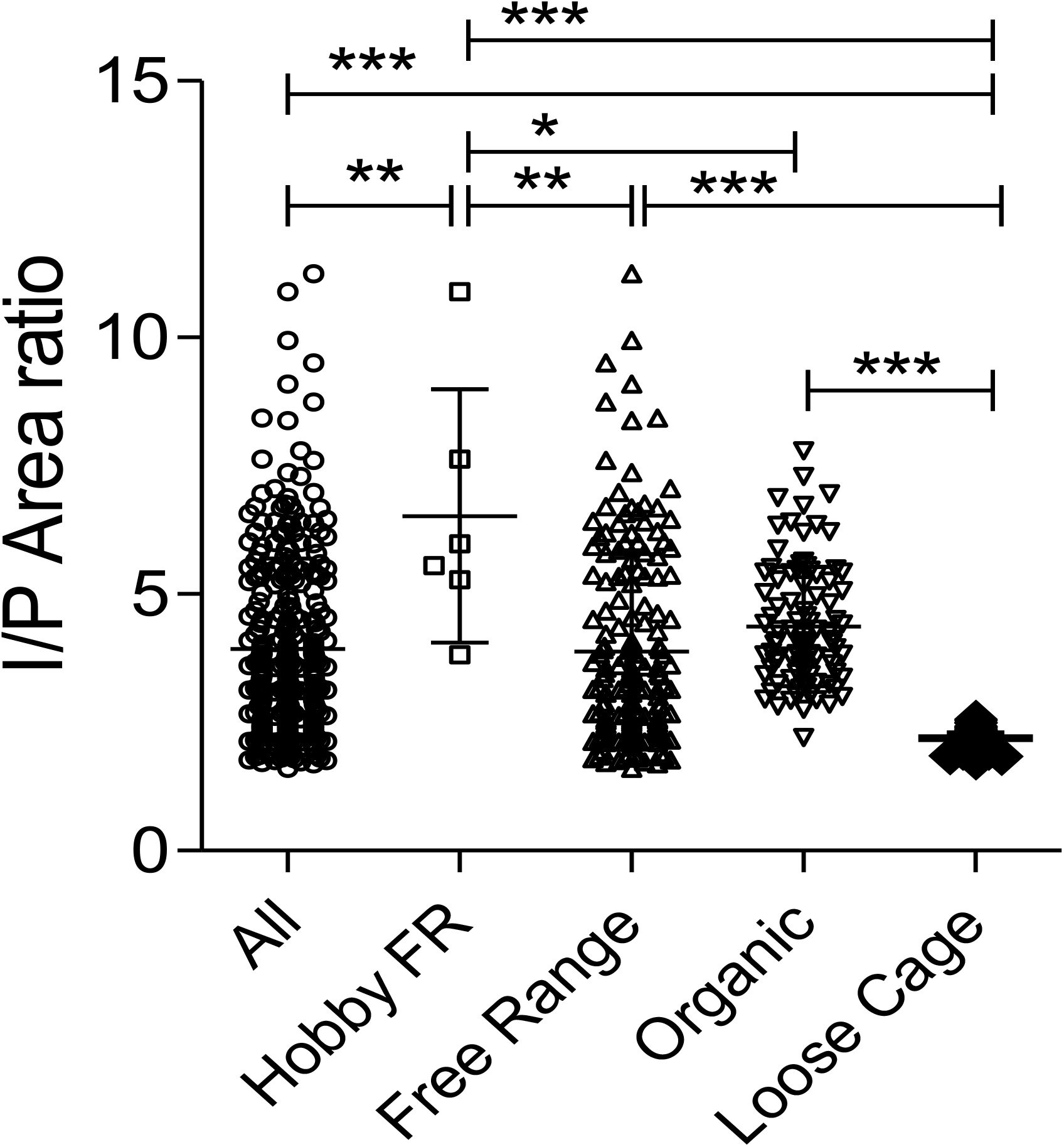
Iris-Pupil (I/P) area ratio of layer birds reared under different commercial systems, All (n = 296), hobby free-range (FR) (n = 6), free-range (n = 169), organic EU-compliant (n = 94), and loose cage housing (n = 27). Individual IP ratio values along with mean ± standard deviation values for each group are presented. Data was analysed by One way ANOVA followed by Bonferroni’s Multiple Comparison Test to assess the significance between two groups. * p<0.05, ** p<0.01, and *** p<0.001.

We next examined whether the housing location had any influence on the IP ratio in laying hens. Four distinct environmental settings were included in the study, i.e., outdoor areas, indoor housing, scratch (enriched litter) system, and indoor pens. The IP ratio of birds reared outdoors was 140.52%, 63.42% and 153.80% greater than the IP ratio of birds in indoor, scratch and pen system respectively. Among these, birds reared in outdoor environments exhibited the highest IP ratio, followed by those housed in the scratch system. In contrast, birds kept in indoor settings or pens demonstrated the lowest IP ratios (Figure 3A). The IP ratios of birds in indoor settings and pens was not statistically (p > 0.05) different (Figure 3A). Statistical analysis suggested that the IP ratio of birds in the outdoor environment was significantly greater (p < 0.001) than that of birds in the scratch, indoor, and pen systems. Furthermore, birds in the scratch system also showed significantly higher IP ratios (p < 0.001) compared to those housed indoors or in pens, suggesting a graded effect of environmental complexity and access to natural stimuli on physiological stress indicators.

**Figure 3.**
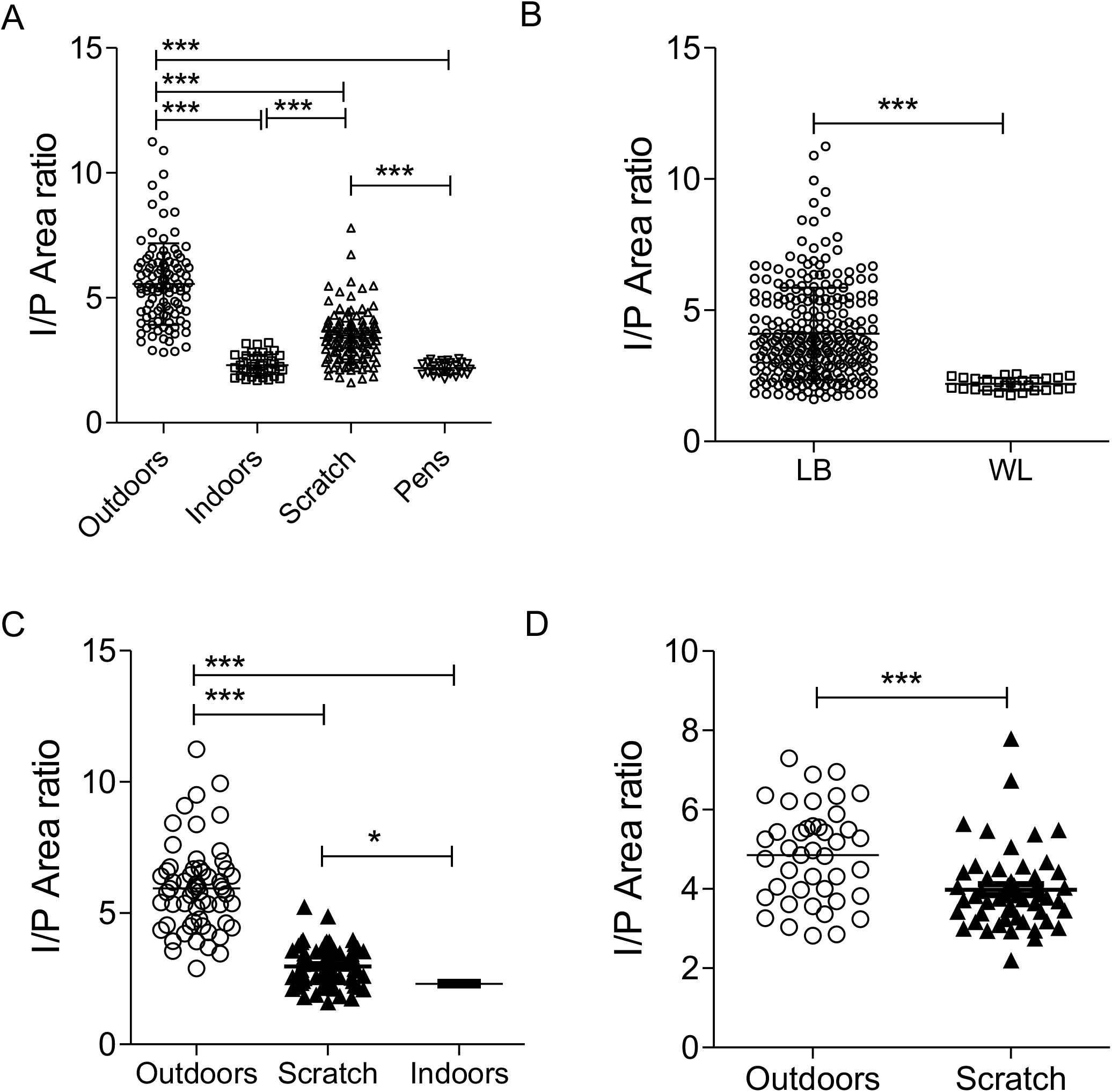
A: Iris-Pupil (I/P) area ratio of layer birds reared under different husbandry systems. Outdoor (n = 60), indoor (n = 38), scratch area (n = 71) and pens (n = 27). Individual IP ratio values along with mean ± standard deviation values for each group are presented in all graphs. Data was analysed by one way ANOVA followed by Bonferroni’s Multiple Comparison Test to assess the significance between two groups. *** p<0.001. B: I/P area ratio of Lohmann Brown and White Leghorn breeds of layers. Data was analysed by student t-test (*** p<0.001). I/P area ratio of layer birds reared under different husbandry systems in Kelly (C) and Olive (D) farm. Data from Kelly farm was compared using one way ANOVA while data from Olive farm was compared using student t-test (* p<0.05 and *** p<0.001).

Two commercial laying hen breeds were included in this study: Lohmann Brown and White Leghorn. We assessed whether the IP ratio varied between breeds by comparing all Lohmann Brown birds (n = 269) with all White Leghorn birds (n = 27). The results showed that Lohmann Brown hens had a significantly higher IP ratio (p < 0.001) than White Leghorn hens (Figure 3B). However, this apparent breed difference is likely confounded by the housing conditions. All White Leghorn hens were housed exclusively in the pen system, while Lohmann Brown hens were represented across all four housing environments (outdoor, indoor, scratch, and pen). When comparing IP ratios between Lohmann Brown hens in the indoor system and White Leghorn hens in the pen system, no significant (p > 0.05) difference was observed (Figure 3A), indicating that the lower IP ratio in White Leghorns is more reflective of the housing system than of a breed-specific trait. Hence, it is reasonable to conclude that breed did not independently influence the IP ratio in this study; rather, environmental conditions were the primary driver of the observed differences. To further validate the influence of the housing environment on the IP ratio, we conducted a subgroup analysis using birds from two farms, i.e., Kelly and Olive farms. These farms provided an opportunity to examine IP ratio differences within a consistent management system while varying only the immediate environment, namely, outdoor, indoor, and scratch areas. At Kelly Farm, which operated under a free-range system, birds were housed in three distinct environments: outdoor, indoor, and scratch. Analysis revealed that birds kept outdoors had a significantly higher IP ratio compared to those housed in both the scratch and indoor systems (p < 0.001; Figure 3C). Additionally, birds in the scratch system showed a significantly higher IP ratio than those kept indoors (p < 0.05; Figure 3C). These findings suggest a clear gradient in IP ratio values corresponding to the degree of environmental enrichment and access to natural conditions, with outdoor environments associated with the highest IP ratios.

A similar pattern was observed at Olive farm, which followed an organic, EU-compliant system and housed birds in both outdoor and scratch environments. Although Olive farm did not include an indoor-only group, comparison between the two available environments again showed a significantly higher IP ratio in birds kept outdoors compared to those in the scratch system (p < 0.001; Figure 3D). The consistency of these results across two separate farms reinforces that housing environment, particularly access to outdoor conditions, has a measurable influence on the physiological state of laying hens as reflected by their IP ratio. These findings support the utility of the IP ratio as a non-invasive, objective welfare indicator sensitive to environmental quality.

## Discussion

This study investigated the use of the iris-to-pupil (IP) ratio as a non-invasive indicator of physiological stress and welfare in laying hens housed under different commercial systems and environmental conditions. Our findings demonstrate that the IP ratio is significantly influenced by the birds’ housing environment, with birds reared in outdoor and enriched settings consistently exhibiting higher IP ratios than those housed indoors or in conventional pen systems. These results are consistent with the hypothesis that more naturalistic, enriched environments reduce stress and promote better welfare outcomes in poultry [24-27]. The IP ratio has been previously validated as a reliable, non-invasive measure of autonomic nervous system (ANS) balance in other species, including primates and cattle [7,8,28,29]. In these studies, elevated IP ratios were associated with reduced sympathetic activation and an overall shift toward optimal autonomic balance indicative of improved emotional and physiological states. Our present findings in laying hens align with this pattern, i.e., birds kept in outdoor or enriched “scratch” environments, which offer greater space, foraging opportunities, and environmental stimuli, showed higher IP ratios, suggesting a state of lower physiological stress and favorable welfare status.

The ANS plays a central role in mediating physiological responses to stress in birds, just as it does in mammals [30-32]. Activation of the sympathetic nervous system leads to pupil dilation, increased heart rate, and mobilization of energy reserves, while parasympathetic dominance supports rest, digestion, and recovery [7,8,33]. As the pupil diameter reflects the dynamic balance between sympathetic and parasympathetic inputs of the ANS, the IP ratio emerges as a quantifiable proxy for this neurophysiological state [34-36]. Under increased sympathetic activation, the pupil dilates, decreasing the IP ratio, whereas parasympathetic dominance leads to pupil constriction, increasing the IP ratio. However, because pupil size is also responsive to changes in ambient light, dilating in low light and constricting in bright light, it is essential that IP ratio measurements are conducted under controlled or consistently measured lighting conditions to ensure that observed differences reflect physiological stress responses rather than lighting artifacts [7,37]. In our study, light intensity was recorded at each site using a standardized method, and correlation analyses demonstrated only a weak relationship between light intensity and IP ratio. Importantly, we observed only weak correlations between IP ratio and both ambient light intensity (r^2^ = 0.25) and birds’ age (r^2^ = 0.05), despite the wide ranges in light intensity (21 to 20,000 lux) and age (25 to 104 weeks). This finding validates the reliability of the IP ratio as a field-applicable welfare metric, demonstrating minimal interference from commonly varying environmental factors. This is especially relevant in field settings, where lighting conditions are often difficult to standardize, yet physiological measurements must remain accurate. This further supports the reliability of the IP ratio as an indicator of autonomic balance, if variations in lighting are accounted for or minimized during data collection.

In our study, the robust differences in IP ratios across systems with differing levels of environmental complexity, strongly suggest that the IP ratio is sensitive to the underlying emotional and physiological states of the birds. The analysis across different production systems further highlighted meaningful welfare differences. Birds in the hobby free-range system showed the highest average IP ratio (6.52 ± 2.47), significantly greater than all other groups, while those in the loose cage system showed the lowest (2.9 ± 0.24), significantly below the cohort average (3.93 ± 1.76). This pattern suggests that extensive outdoor access and reduced confinement contribute to better welfare states, while restrictive, barren environments are associated with increased physiological stress. These findings are consistent with existing literature, which has shown that cage-free and outdoor systems are associated with improved behavioral expression and lower indicators of stress in poultry [38-40]. Birds housed in the hobby free-range system exhibited markedly higher IP ratio compared to those in other commercial housing environments. Specifically, the IP ratio in the hobby free-range group was 68.23% higher than in birds from standard free-range systems, 49.22% higher than those housed in organic systems, and 198.16% greater than in birds from loose cage systems. These substantial differences suggest the adaptations in autonomic balance, and potentially lower stress levels, experienced by birds in the hobby free-range environment, which typically provides more enriched and less densely populated outdoor conditions. Importantly, these differences in IP ratio were disproportionate to the variability observed in ambient light intensity across the systems, suggesting that the elevated ratios in the hobby free-range birds are not simply an artifact of lighting differences. This is further supported by the weak correlation found between light intensity and IP ratio mentioned before. Hence, these findings reinforce the role of environmental complexity and management practices in shaping physiological welfare outcomes in laying hens and highlight the sensitivity of the IP ratio as a reliable indicator of these welfare changes.

The data from this study revealed striking differences in the IP ratio between birds reared in different housing environments, with the highest values observed in birds reared outdoors. Specifically, the IP ratio of birds in the outdoor system was 140.52% greater than those housed indoors, 63.42% greater than birds in the scratch system, and 153.80% greater than birds housed in indoor pens. These significant elevations in IP ratio among outdoor-reared birds suggest that access to natural light, fresh air, and greater environmental complexity plays a critical role in reducing physiological stress, as reflected in improved autonomic balance. The outdoor environment provides birds with a richer array of sensory, spatial, and behavioral stimuli, all of which are known to support better welfare outcomes. Natural settings facilitate expression of species-specific behaviors such as foraging, dust bathing, and exploration activities which are often limited or absent in more confined housing systems. The elevated IP ratios observed in these birds indicate a shift toward a more balanced autonomic nervous system state, potentially characterized by reduced sympathetic tone and increased parasympathetic influence, both of which are consistent with reduced stress and improved welfare. In contrast, birds housed indoors or in pens, which represent more confined and stimulus-poor environments, showed the lowest IP ratios, indicative of a higher baseline stress load or reduced environmental engagement. The scratch system, although better than indoor pens due to the presence of litter and some enrichment, still fell significantly short of the benefits observed in fully outdoor systems. Again, these observed differences in IP ratio were far greater than could be accounted for by variation in ambient light intensity alone, which showed only a weak correlation with IP values. This strengthens the interpretation that environmental quality, specifically access to outdoor space is a key determinant of physiological welfare status in laying hens. These findings align with existing literature highlighting the benefits of outdoor access on behavioral health [38-40] and support the inclusion of IP ratio as a robust, quantifiable tool for evaluating animal welfare across production systems.

To further refine our understanding, we conducted within-farm comparisons at Kelly and Olive farms. In both cases, birds housed outdoors exhibited significantly higher IP ratios than those in scratch or indoor environments, despite being managed under the same general system (free-range and organic, respectively). These results highlight the value of outdoor access itself, rather than the broader management label, as a critical determinant of welfare. Such findings add depth to ongoing debates about the relative value of “free-range” and “organic” labeling, suggesting that detailed assessment of specific environmental components may be more informative than broad system classification. Breed differences in IP ratio were initially observed, with Lohmann Brown birds showing significantly higher IP ratios than White Leghorns. However, this difference was attributable to housing conditions, as all White Leghorns were housed in pens, while Lohmann Browns were distributed across all systems. Direct comparison of Lohmann Browns housed indoors and White Leghorns in pens revealed no significant differences, confirming that environmental factors, rather than breed, primarily influenced the IP ratio in this study.

Pupillometry, and specifically the use of the IP ratio, offers several compelling advantages for welfare assessment in laying hens. As demonstrated in this study, the IP ratio effectively discriminates between birds housed in different environmental conditions, with higher IP ratios observed in birds reared in more enriched, outdoor systems and lower ratios in confined or barren environments such as indoor pens. This sensitivity to housing quality merits the IP ratio value as a proxy for ANS balance, which is a critical physiological correlate of stress/welfare. One of the most significant benefits of using the IP ratio is its non-invasive nature, pupillary measurements can be obtained through simple photography, thereby eliminating the need for physical restraint or handling of animals. This not only avoids introducing unnecessary stress during data collection, which can confound the physiological parameters being measured, but also aligns with ethical standards for humane research practices [41-43]. Furthermore, pupillometry is highly cost-effective. Unlike many traditional physiological assessments that require invasive sampling or specialized laboratory equipment,[44,45] this method only requires a digital camera and open-source image analysis software, making it accessible for both research and on-farm monitoring.

Taken together, these findings strongly support the utility of the IP ratio as an objective, non-invasive indicator of welfare in laying hens, capable of detecting meaningful differences across housing environments. By reflecting the balance of the autonomic nervous system, this simple measure provides insight into the birds’ internal physiological state without the need for invasive procedures or complex equipment. Given its sensitivity, field applicability, and biological relevance, the IP ratio holds significant promise for integration into future welfare assessment protocols in poultry and potentially other avian species. Tools such as the IP ratio not only offer a non-invasive and objective measure of physiological stress but also represent a pathway toward technology-based solutions for welfare monitoring. With advancements in artificial intelligence and digital imaging, IP ratio measurements can be automated using high-resolution imaging and machine learning algorithms, enabling large-scale, real-time welfare assessment with minimal human intervention. Such integration of digital technologies can enhance the accuracy, efficiency, and scalability of welfare evaluations, making them more accessible to both commercial operations and research settings. As the demand for transparent, science-based welfare standards continues to grow,[46,47] adopting data-driven approaches like the IP ratio can play a pivotal role in modernizing and standardizing welfare monitoring in the poultry industry.

In conclusion, this study provides strong evidence supporting the use of the IP ratio as a reliable, non-invasive, and field-applicable indicator of welfare in laying hens. By demonstrating consistent and significant differences in IP ratio across various housing environments, independent of ambient light intensity and age, we establish the physiological sensitivity of this measure to environmental quality and stress exposure. The minimal equipment requirements, ease of data collection through photography, and compatibility with digital imaging and AI technologies further enhance the utility of this method for practical on-farm welfare monitoring. As global standards for animal welfare continue to evolve, incorporating science-based, objective tools such as the IP ratio can offer producers and researchers a meaningful, cost-effective means of assessing and improving the lives of commercial poultry. These findings lay the groundwork for broader application of pupillometry across avian species and production systems, and future work should aim to standardize protocols and validate the IP ratio against behavioral and endocrine markers of stress to strengthen its role in welfare assessments.

